# Stochastic Epigenetic Mutations: Reliable Detection and Associations with Cardiovascular Aging

**DOI:** 10.1101/2023.12.12.571149

**Authors:** Yaroslav Markov, Morgan Levine, Albert T. Higgins-Chen

## Abstract

Stochastic Epigenetic Mutations (SEMs) have been proposed as novel aging biomarkers that have the potential to capture heterogeneity in age-related DNA methylation (DNAme) changes. SEMs are defined as outlier methylation patterns at cytosine-guanine dinucleotide (CpG) sites, categorized as hypermethylated (hyperSEM) or hypomethylated (hypoSEM) relative to a reference. While individual SEMs are rarely consistent across subjects, the SEM load – the total number of SEMs – increases with age. However, given poor technical reliability of measurement for many DNA methylation sites, we posited that many outliers might represent technical noise. Our study of whole blood samples from 36 individuals, each measured twice, found that 23.3% of hypoSEM and 45.6% hyperSEM are not shared between replicates. This diminishes the reliability of SEM loads, where intraclass correlation coefficients are 0.96 for hypoSEM and 0.90 for hyperSEM. We linked SEM reliability to multiple factors, including blood cell type composition, probe beta-value statistics, and presence of SNPs. A machine learning approach, leveraging these factors, filtered unreliable SEMs, enhancing reliability in a separate dataset of technical replicates from 128 individuals. Analysis of the Framingham Heart Study confirmed previously reported SEM association with mortality and revealed novel connections to cardiovascular disease. We discover that associations with aging outcomes are primarily driven by hypoSEMs at baseline methylated probes and hyperSEMs at baseline unmethylated probes, which are the same subsets that demonstrate highest technical reliability. These aging associations are preserved after filtering out unreliable SEMs and are enhanced after adjusting for blood cell composition. Finally, we utilize these insights to formulate best practices for SEM detection and introduce a novel R package, *SEMdetectR*, which utilizes parallel programming for efficient SEM detection with comprehensive options for detection, filtering, and analysis.

## INTRODUCTION

Many age-related changes are universal or occur in most individuals, but certain pertinent changes may manifest in only a few individuals. For instance, a specific somatic mutation might arise in a limited subset of people, even though the collective burden of somatic mutations increases with age throughout the population <u>[1]</u>. Epigenetic changes are widespread in aging and disease <u>[2]</u>, and many studies have associated aging with shifts in DNA methylation (DNAme) at cytosine-guanine dinucleotides (CpGs), an epigenetic alteration that modulates gene transcription <u>[3]</u>. Epigenetic clocks, based on DNAme, are frequently employed as aging biomarkers and have displayed associations with a multitude of aging outcomes and risk determinants <u>[4,5]</u>. Most existing studies, such as epigenome-wide association or epigenetic clock studies, identify CpGs that either undergo hypo- or hyper-methylation with age or those that change in variance; detecting such changes necessitates their prevalence in a significant fraction of the population.

Stochastic Epigenetic Mutations (SEMs) have been proposed as a complementary DNAme-based biomarker of aging. Conceptually, a SEM is an outlier in DNAme value at a specific genomic site when compared with all subjects in a given population or cohort. Inherently, specific SEM sites are seldom consistent across individuals, often being unique to one person within a cohort. Even the most recurrent SEMs manifest in fewer than 1% of subjects <u>[6,7]</u>. Nonetheless, as with somatic mutations, SEM load (the total burden of SEMs in an individual) increases with age across the population as well as longitudinally within individuals during aging <u>[6,8–10]</u>. Gentilini et al. defined SEM as extreme DNAme outliers that exceed three times the interquartile ranges (IQR) from the first quartile (Q1-(3IQR)) or the third quartile (Q3+(3IQR)) of the cohort of interest <u>[7]</u>. SEM load is then typically quantified by simply counting the number of outliers observed in an individual and presented on a logarithmic scale (i.e., log10(SEM count)) due to the exponential increase in SEM with age. SEM load can vary by sex, genetic ancestry, and blood cell composition, though these do not substantially modify the effect of age <u>[6,8,11]</u>. Smoking, obesity, limited education, and environmental exposures may augment the effect of aging on SEM load <u>[8,10,11]</u>. Total SEM load has been associated with future risk of cancer <u>[9]</u> and outcomes in Parkinson’s disease including mortality <u>[12]</u>, though another study found only a nominal association with mortality risk and no cross-sectional associations with a variety of age-related diseases <u>[8]</u>. Though both SEM load and epigenetic clock values increase with age, they are weakly correlated, and the association of SEM with various cancers remains largely unchanged after adjusting for epigenetic clock values, suggesting they might offer complementary information <u>[9,13]</u>.

SEM load and individual SEMs remain relatively under-researched in aging when compared to epigenetic clocks or individual CpGs that undergo hyper- and hypo-methylation with age. One limiting factor is the absence of readily accessible, publicly available code – outlined in any publication – for detecting SEMs or calculating SEM load. This stands in contrast to epigenetic clocks for which multiple published resources exist <u>[14–18]</u>.

There also remain several unresolved questions surrounding the optimal definition of SEMs and SEM load. For instance, it is uncertain whether all SEMs are best grouped into a single metric, categorized into two categories as hypo- and hyper-methylated outliers (hypo- and hyperSEMs respectively), or binned into more detailed SEM subtypes. Currently, SEMs are categorized into hypo- and hyperSEMs; for example, a longitudinal twin study indicated that hyperSEM burden has a stronger association with B cell composition, genetic factors, and current cancer diagnosis in blood than hypoSEM <u>[6]</u>. However, the potential benefits of more detailed categories, or other variations in the definition of SEMs, remain to be seen. In particular, it is unknown if different types of SEMs may have different associations, or if specific subtypes of SEM are responsible for the observed associations with cancer risk or mortality.

Another concern is the technical reliability of SEM detection. Given that each SEM, by its definition, appears in a limited number of individuals, it becomes challenging to ascertain if the SEM represents a genuine biological outlier or merely stems from measurement errors. In fact, poor sample quality, as indicated by CpG detection P-values, correlates with an increased SEM burden <u>[6]</u>. DNAme at specific sites is known to exhibit substantial technical variation, which can sometimes overshadow regular biological variations, leading to decreased test-retest reliability <u>[19,20]</u>. For example, up to 77.5% of 450K probes show poor reliability (intraclass correlation coefficient ICC < 0.5), especially for low-variance CpGs <u>[21]</u>. Importantly, this unreliability was not connected to the heteroscedasticity of CpG beta-values - in fact, DNAme M-values present even lower ICC values <u>[19,22]</u>. When assessing SEM stability over time, findings suggest that SEMs remain consistent in only about 70% of instances <u>[9]</u>. Some of this inconsistency might arise from technical noise rather than genuine intraindividual biological changes. SEM loads from 15 technical replicates from a single sample correlate with each other in the range of 0.8-0.92 <u>[10]</u>. However, it is unknown how technical variance in SEM load compares to biological variance across multiple samples. Furthermore, the reliability of individual SEMs has yet to be examined.

Here, we systematically investigate the reliability of both individual SEMs and SEM loads, as well as their associations with age, mortality, and age-related cardiovascular phenotypes. We test the hypothesis that a substantial proportion of SEMs represent technical noise, characterize the features that predict whether a SEM is unreliable, and identify SEM subtypes that drive associations with mortality and cardiovascular disease. We formalize our findings in terms of a set of best practices for SEM detection, as well as *SEMdetectR*, a publicly available R package for SEM detection and characterization.

## RESULTS

We hypothesized that many SEMs could represent measurement noise rather than genuine biological outliers, which would imply that they would not be shared between technical replicates. To investigate the influence of probe and sample characteristics on SEM reliability, we analyzed the publicly available GSE55763 dataset of whole blood samples, where our analyses centered on 36 pairs of technical replicates, using the other 2,664 samples from the dataset as a reference. The large reference dataset is important for calling SEMs which are by definition outliers relative to a reference distribution, and for deriving probe statistics to investigate determinants of reliability.

### Assessing SEM Reliability with the Standard IQR-Based Method

As expected, SEM loads were correlated with age in the 36 pairs of replicates (log10 hypoSEM r = 0.28, p = 0.016; log10 hyperSEM r = 0.38, p = 8.3e-4) (**Figure 1a**). Mean SEM loads were approximately 705 hypoSEM and 1,118 hyperSEM, while median SEM loads were 460 for hypoSEM and 380 for hyperSEM, reflecting the skewed distribution and exponential accumulation of SEMs with aging. Hypo- and hyperSEM loads were strongly correlated with each other (r = 0.88; p < 2.2e-16) (**Supplementary Figure 1**). There were no statistically significant differences in total SEM loads between batches of replicates (hypoSEMs p = 0.583; hyperSEMs p = 0.067), and SEM loads were correlated between replicates (hypoSEM r = 0.97, hyperSEM r = 0.90) (**Figure 1b**).

**FIGURE 1.**
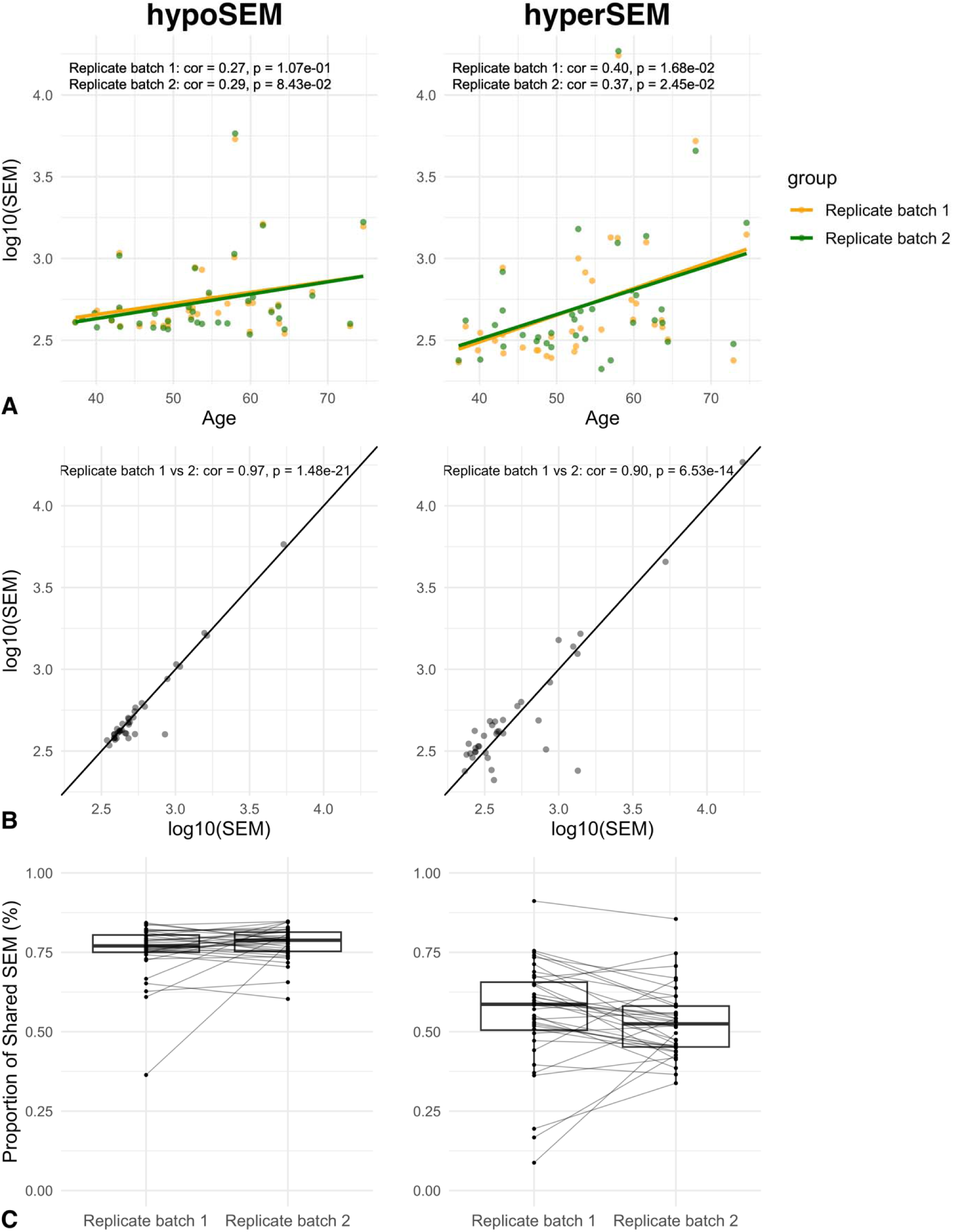
Age associations and technical reliability of SEM in whole blood. Figure 1a: Scatterplot illustrating the relationship between age and hypoSEM (left) and hyperSEM (right) loads in the GSE55763 dataset. The two batches of replicates are color-coded, with one batch represented in yellow and the other in green. Figure 1b: Scatterplot illustrating the agreement between hypoSEM (left) and hyperSEM (right) loads across the two batches of replicates in GSE55763 dataset. The diagonal lines represent the ideal scenario of perfect agreement (slope = 1). Figure 1c: Boxplots representing the proportion of shared hypoSEMs (left) and hyperSEMs (right) out of the total number of hypo- and hyperSEMs, respectively, for each sample in the GSE55763 dataset. Different boxplots illustrate the technical replicates, with lines connecting replicates from the same subjects.

We examined the concordance of individual SEMs between technical replicates and found that the mean percent of shared SEM was 76% for hypoSEM and 54% for hyperSEM, suggesting worse reliability for hyperSEM (**Figure 1c**). However, proportions of shared SEM exhibited considerable variation across subjects – spanning roughly 36-85% for hypoSEM and 9-92% for hyperSEM – and were not significantly different between replicate batches (hypoSEM p = 0.134; hyperSEM p = 0.209).

These results raise concerns about the reliability of individual SEMs, which could potentially impact downstream analyses including calculation of SEM loads. Intraclass correlation coefficient (ICC) values for SEM loads were approximately 0.96 (95% CI: 0.93 < ICC < 0.98) for hypoSEM and 0.90 (95% CI: 0.81 < ICC < 0.95) for hyperSEM. When adjusted for age, the ICC scores remained at 0.96 (95% CI: 0.92 < ICC < 0.98) for hypoSEM and decreased slightly to 0.88 (95% CI: 0.78 < ICC < 0.94) for hyperSEM.

### Identifying Drivers of SEM Concordance

We aimed to better understand why SEMs are inconsistently shared across replicates and identify ways to improve SEM reliability. First, for all SEMs that were detected in at least one replicate (for any of the 36 samples), we recorded the beta value of the corresponding CpG in the second replicate. Using this information, we plotted the “deltaIQR” of the second replicates – representing the number of interquartile ranges the beta value is below Q1 for hypoSEM or above Q3 for hyperSEM – and observed a bimodal distribution (**Figure 2**). In the first mode, the second replicates were either classified as a SEM, or were directionally consistent with SEMs from corresponding first replicates, albeit not reaching the 3 x deltaIQR threshold. In contrast, the second mode had the second replicates considerably distanced from the 3 x deltaIQR mark, with some even exceeding Q1 for hypoSEM or dropping below Q3 for hyperSEM. We noted the boundary between these two modes was approximately at a 1.5 x deltaIQR cutoff, formally corroborated by gaussian mixture modeling analysis (cutoff 1.26 for hypoSEM, 1.49 for hyperSEM). Stemming from these observations, we classified SEMs into three categories for further analysis:

1. shared SEMs: both replicates surpassed the 3-IQR threshold (57.9% for hypoSEM and 55.3% for hyperSEM);
2. almost-shared SEMs: one replicate surpassed the 3-IQR threshold, and the other surpassed the 1.5-IQR threshold (26.3% for hypoSEM and 24.5% for hyperSEM);
3. unshared SEMs: one replicate surpassed the 3-IQR threshold, and the other did not reach the 1.5-IQR threshold (15.7% for hypoSEM and 20.2% for hyperSEM).

**FIGURE 2.**
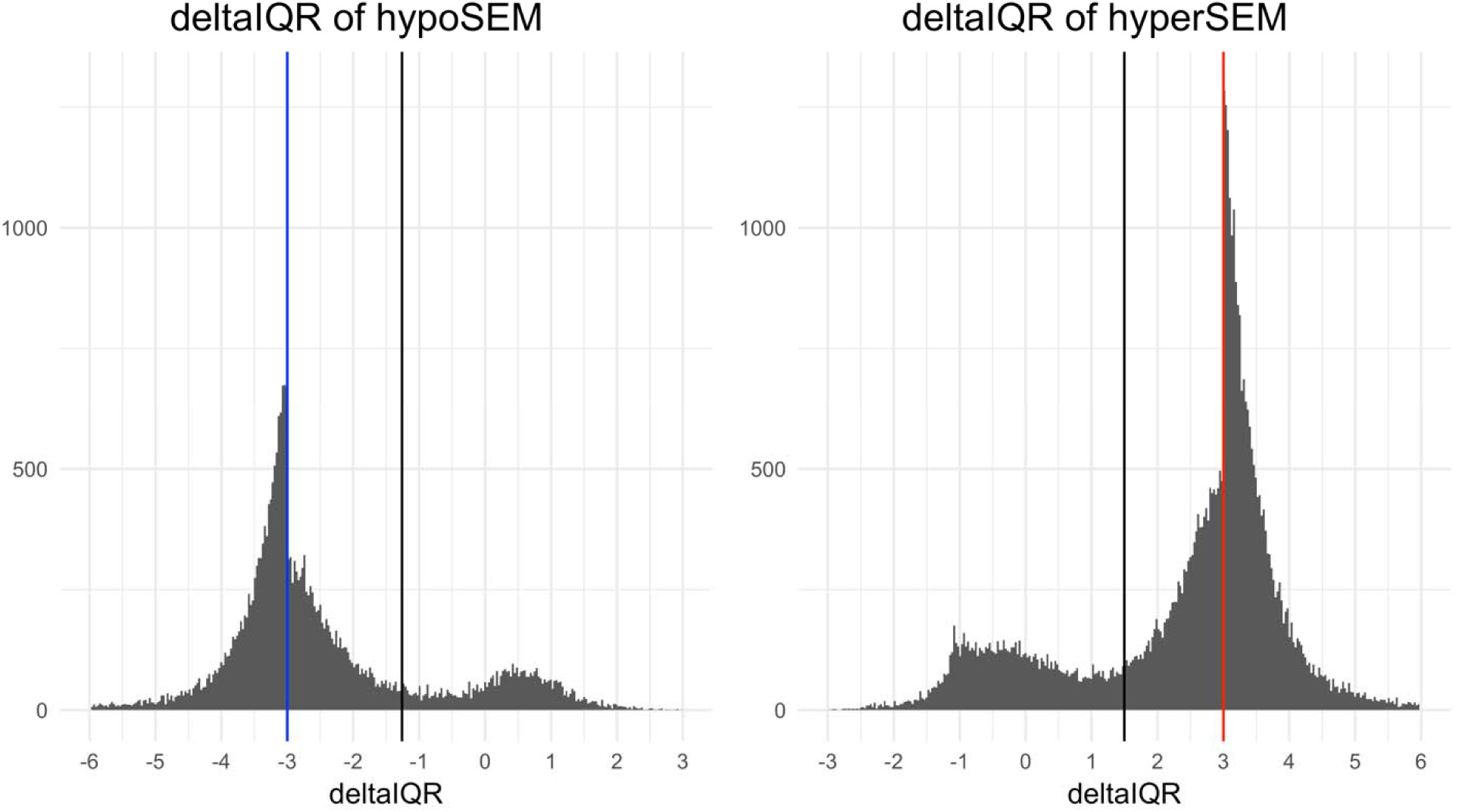
Bimodal deltaIQR distributions. Histograms illustrating the distribution of ‘deltaIQR’ values in the GSE55763 dataset for hypoSEMs (left panel) and hyperSEMs (right panel). For each SEM detected in at least one replicate, we plot the deltaIQR for the other technical replicate. The ‘deltaIQR’ is calculated as the number of interquartile ranges (IQRs) below the first quartile (Q1) for hypoSEMs or above the third quartile (Q3) for hyperSEMs regarding the beta value of the second technical replicate. The blue and red vertical lines denote the original 3 x IQR threshold utilized for SEM detection in hypo- and hyperSEMs, respectively. Black vertical lines represent the points of intersection between the two distributions as modeled by Gaussian mixture modeling. For clarity of presentation, outliers residing in the extended tails of these distributions are omitted.

Next, we systematically analyzed the three SEM categories in relation to probe and sample characteristics. Results concerning all probe and sample characteristics investigated can be found in **Supplementary Figures 2** and **3**, while particularly notable results are shown in **Figure 3**. We discovered significant differences in probe statistics, such as mean, IQR, quantiles, and others, between shared and unshared SEMs (**Figure 3** and **Supplementary Figure 2**). In particular, unshared SEMs had lower IQR and standard deviation compared to shared and almost shared SEMs (p-value < 2.2e-16 for both comparisons, for both hyper- and hypoSEMs), consistent with previously reported lower reliability for CpGs with low variance <u>[20]</u>. In contrast, the differences between shared and almost shared SEMs were not uniformly significant across hypo- and hyperSEMs; there was a difference in standard deviation for hypoSEMs and IQR for hyperSEMs (p-value < 2.2e-16 for both). For sample characteristics, we observed that both hypoSEM and hyperSEM were more likely to be shared between replicates in samples estimated to have elevated levels of CD8T and B cells (p < 2.2e-16 for both) and reduced granulocyte levels (p < 2.2e-16) (**Figure 3**).

**FIGURE 3.**
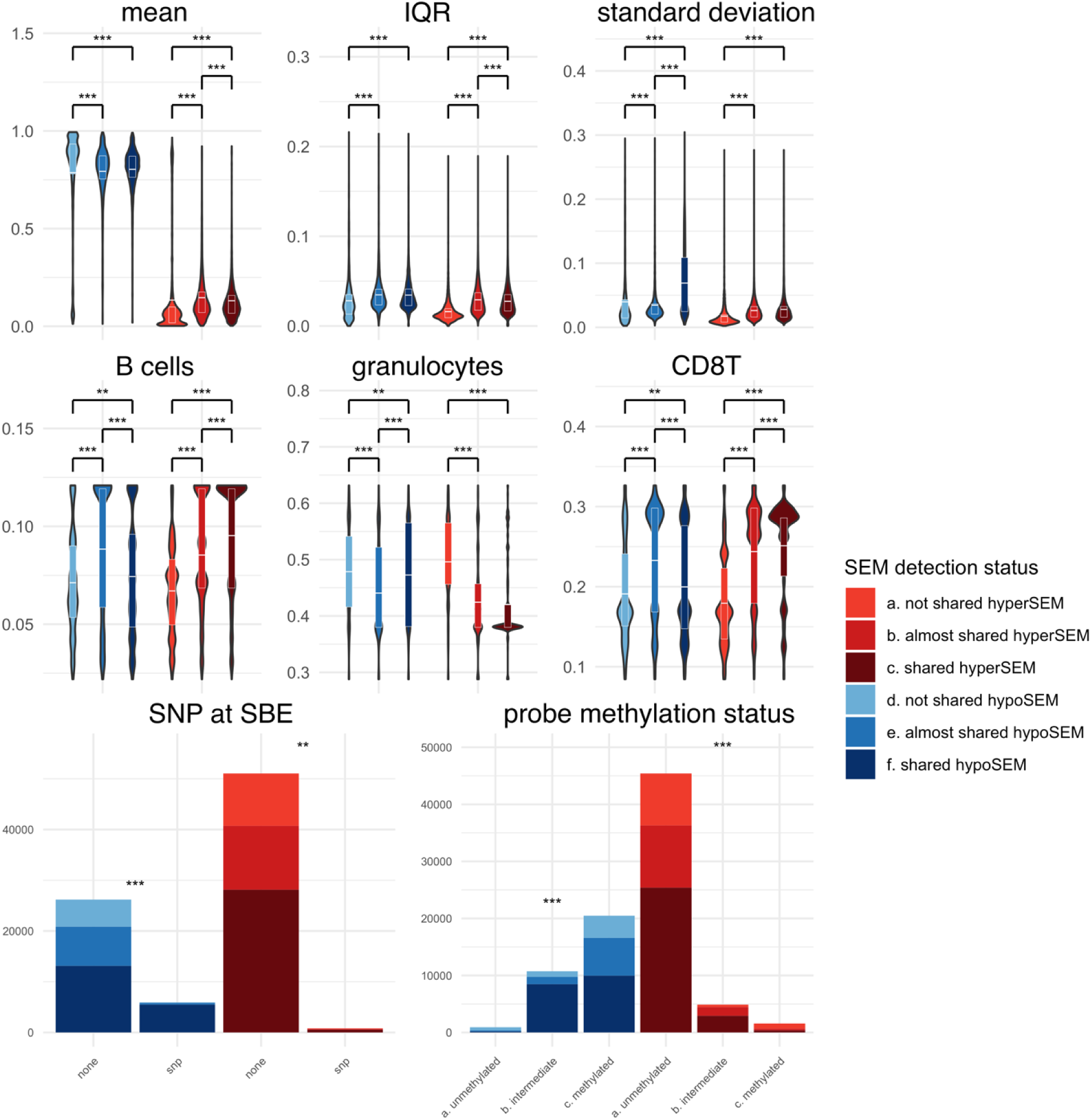
Selected features analyzed in relation to the reliability status of SEMs. hypoSEMs illustrated in blue and hyperSEMs in red. Continuous features are displayed through violin plots, while categorical features are displayed through stacked bar plots. For mean, IQR, and standard deviation, units are in terms of methylation beta-values. For B cells, granulocytes, and CD8T cells, unit are in terms of the proportion of cells (0 to 1). For SNPs at SBE and probe methylation status, y-axis corresponds to the total counts of SEMs. Statistically significant differences between violin plots are indicated by asterisks, with white horizontal lines representing the mean values and quartiles. For the stacked barplots, asterisks placed atop denote statistically significant differences within hypo- or hyperSEM groups. Significance values were calculated by the Chi-squared test for categorical variables and the Mann-Whitney test for continuous variables, and are denoted as follows: *** for p < 2e-16 and ** for p < 3.8e-4 (Bonferroni corrected significance value).

The presence of Single Nucleotide Polymorphisms (SNPs) overlapping with the single base extension site (SBE), CpG, or present elsewhere in the probe was linked to an increase in hypoSEM reliability (p < 2.2e-16 for all) (**Figure 3** and **Supplementary Figure 3**). This association was weaker, albeit still statistically significant, for hyperSEM reliability (probe: p = 1.9e-13; CpG: p < 2.2e-16; SBE: < 1.8e-10). Probes with SNPs located at SBE had a negligible number of hyperSEMs (SBE: 1.5% of total) but a considerable proportion of hypoSEMs (18.5%). Further investigating SNPs effects on SEM reliability, we found that SEM-containing probes with SNPs at the SBE constituted a minor portion of all SEM-containing probes (∼3.5%), and removing them only slightly reduced the ICC scores from 0.96 to 0.95 for hypoSEM.

### Probes containing hypoSEMs vs probes containing hyperSEMs

We observed that probe statistics not only varied based on SEM reliability but also differed between hypo- and hyperSEMs. For instance, metrics such as mean, sum under the curve, minimum, and quartiles were typically higher for hypoSEMs than for hyperSEMs (**Figure 3** and **Supplementary Figure 2**). Drawing from these observations, we postulated that hypo- and hyperSEMs likely arise from probes exhibiting distinct methylation patterns. To validate this idea, we utilized the full reference cohort to categorize all probes into three groups based on their DNAme beta values: unmethylated, intermediate, and methylated (average means 0.08, 0.50, and 0.84 respectively). Our investigation showed that a substantial majority of hypoSEMs (∼63%) originated from methylated probes, whereas a predominant portion of hyperSEMs (∼87%) came from unmethylated probes.

Notably, the most unreliable SEMs were hypoSEMs originating from unmethylated probes and hyperSEMs from methylated probes (**Figure 3**). This observation was further supported by ICC scores: for hypoSEM from unmethylated probes, the score was 0.78 (95% CI: 0.6 < ICC < 0.88), while for hyperSEMs from methylated probes, it was 0.32 (95% CI: 0 < ICC < 0.59) (**Supplementary Figure 4**). Conversely, hypoSEMs from methylated probes and hyperSEMs from unmethylated probes had much higher ICC scores of 0.94 (95% CI: 0.88 < ICC < 0.96) and 0.9 (95% CI: 0.81 < ICC < 0.94) respectively, closely aligning in reliability with the overall hypo- and hyperSEM loads.

### Addressing SEM Reliability

The original IQR-based approach for SEM detection is designed for normally distributed data, but DNAme beta values typically exhibit pronounced skewness in their distributions across subjects. Initially, we hypothesized that this discrepancy between the SEM detection method’s underlying assumptions and the actual distributional characteristics of DNAme data might be contributing to the observed reliability issues. To investigate this, we utilized a skewness-adjusted algorithm for outlier detection, as described by <u>[23]</u>). Contrary to our expectations, this approach did not improve reliability; in fact, it decreased the percentage of shared SEM and worsened the ICC scores (data not shown). These outcomes prompted us to experiment with machine learning techniques to predict the reliability of SEMs identified by the original IQR-based method.

We trained two separate random forest (RF) classifiers: one for predicting the reliability of hypoSEMs, and the other for hyperSEMs. Essentially, these models predicted whether, for a given SEM, we would identify a SEM in its paired technical replicate. As previously highlighted, no definitive cutoff existed between shared SEMs and almost-shared SEMs in terms of the deltaIQR of the second replicate, and there were only subtle differences in terms of probe and sample characteristics. Thus, we grouped almost-shared SEMs with shared SEMs for the ground truth of the models. It is important to note that this does not make the SEM detection algorithm more lenient; the models still only consider SEMs initially detected at a threshold of at least 3 x deltaIQR, but it requires the second replicates to meet a threshold of 1.5 x deltaIQR.

Following hyperparameter tuning and cross-validation on the training data split, the finalized models yielded AUC scores of approximately 0.93 for hypoSEMs and 0.95 for hyperSEMs when applied to the test data split. As expected from the high AUC values, our RF-based models notably increased proportions of shared SEMs by on average approximately 8% (from 76% to 84%) for hypoSEMs (p < 2.4e-18) and 19% (from 54% to 73%) for hyperSEMs (p < 3.3e−20) (**Figure 4a**). Intriguingly, even though the models were not explicitly trained to enhance ICC scores, they improved these metrics to 0.99 or higher for both hypoSEMs and hyperSEMs, with confidence intervals of 95% CI 0.994 < ICC < 0.998 and 95% CI 0.985 < ICC < 0.996, respectively. Notably, the ICC scores improved across all subtypes of SEMs (e.g., hypoSEMs from methylated probes), as shown in **Figure 4b**. The RF method decreased loads of SEMs per individuals, particularly hypoSEMs in unmethylated probes (removed on average 68%) and hyperSEMs in methylated probes (removed on average 49%). In absolute numbers, however, it preserved ∼71,000 SEM remaining out of initial ∼84,000 (combined for all replicates) (**Figure 4c**).

**FIGURE 4.**
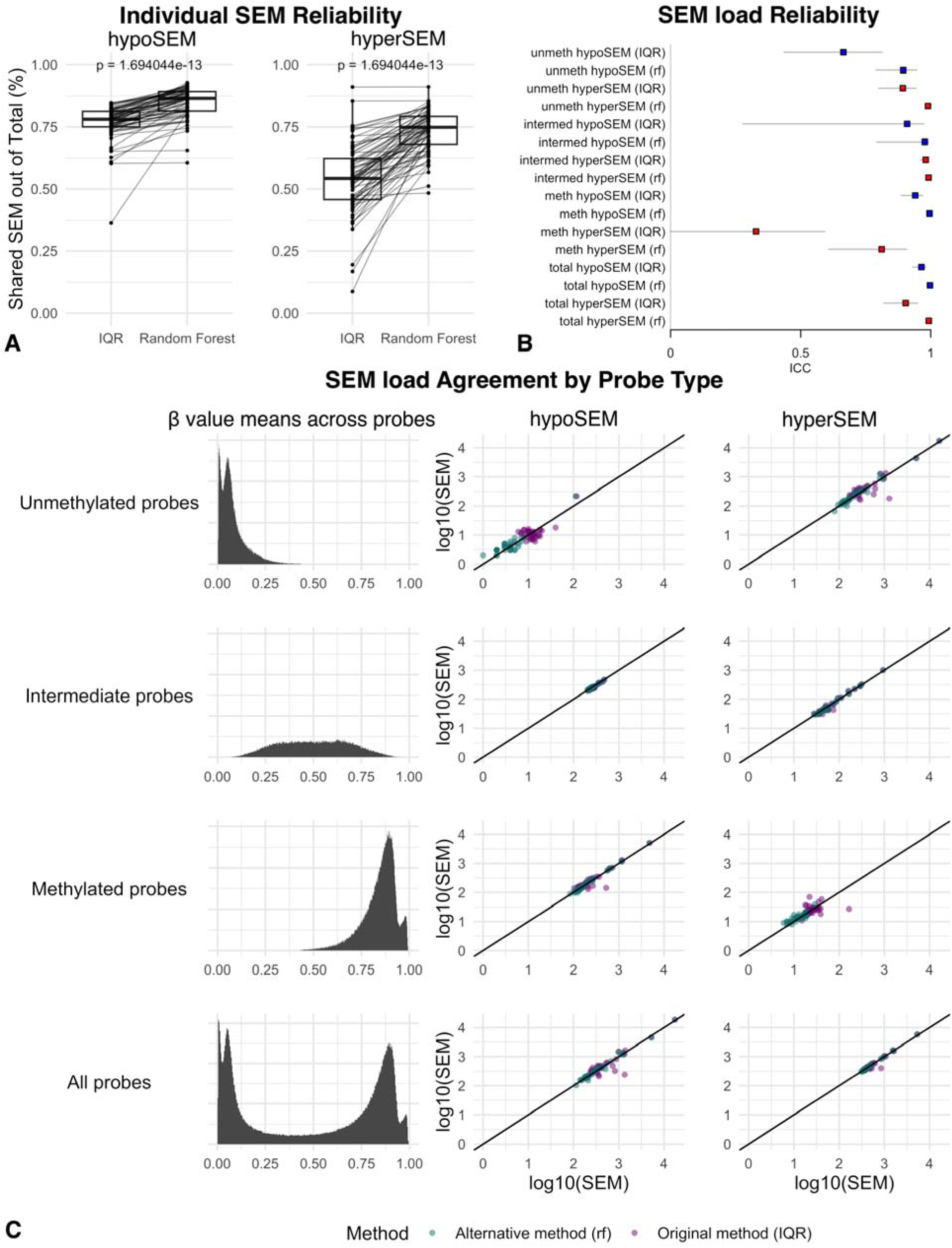
SEM reliability: Effect of the random forest filter in GSE55763. Figure 4a: Boxplots depicting the proportion of shared hypoSEMs (left) and hyperSEMs (right) out of the total number of identified hypo- and hyperSEMs, respectively, as detected by the original IQR-based method vs filtered by the RF-based method in the GSE55763 dataset. Different boxplots represent the different methods used, with lines connecting the same samples analyzed by the two different methods. The replicate batch is not shown in this representation. Figure 4b: This forest plot displays the ICC scores for the SEM load subtypes analyzed by the original IQR-based method vs filtered by the alternative RF-based method within the GSE55763 dataset. The hypoSEM loads are color-coded in blue, while the hyperSEM loads are depicted in red. Horizontal bars denote the lower and upper bounds of the ICC measurements. The label ‘Total’ refers to the SEM loads obtained from all probes. Labels ‘unmeth,’ ‘intermed,’ and ‘meth’ indicate the SEM loads originating from unmethylated, intermediate, and methylated probes, respectively, as classified through reference dataset analysis. The tag (rf) indicates that the SEMs were detected with the RF-based filtering, while the rest were detected using the IQR-based method. Figure 4c: Scatterplots for the SEM subtypes demonstrating the agreement in SEM loads between the first and second batches of replicates in the GSE55763 dataset. SEM loads detected with the original method are illustrated in green, while those identified with the RF-based filtering are depicted in purple.

For the hypoSEM model, the final features, ranked from most to least important according to the mean decrease in Gini coefficient, were: probe IQR, minimum, range, mean, and maximum statistics, deltaIQR of the first replicate, probe standard deviation, B cell, CD8T, and CD4T levels, probe genomic location, number of CpGs overlapping with probe, probe color channel, presence of a SNP at SBE, and the methylation status of the probe. For the hyperSEM model, the ranked features included: CD8T level, probe range, standard deviation, skewness, Q3, and IQR statistics, B cell level, deltaIQR of the first replicate, probe kurtosis, CD4T and NK levels, regulatory features overlapping with probe, probe methylation status, overlap with differentially methylated regions, DNase hypersensitive sites, and enhancers.

To validate our models, we utilized the DNA methylation dataset from the NIEHS Sister Study (GSE174422), which consists of 128 pairs of technical duplicate blood DNA samples from women analyzed on 450K arrays. We identified SEMs using both the original method and our RF-based method, and then compared the results. We again found statistically significant increases in the proportions of shared SEMs — on average by roughly 7% (from 63% to 70%) (p < 1.9e-44) for hypoSEM and 17.8% (from 44.2% to 62%) for hyperSEM (p < 6.1e-69) (**Figure 5a**). ICC scores also improved, though the gains were primarily seen for hyperSEM: ICC for hypoSEM increased from 0.75 (95% CI: 0.66 < ICC < 0.81) to 0.76 (95% CI: 0.68 < ICC < 0.83), while ICC for hyperSEM increased from 0.56 (95% CI: 0.44 < ICC < 0.67) to 0.71 (95% CI: 0.62 < ICC < 0.79) (**Figure 5b**). Similarly to GSE55763, the RF-based method removed most hypoSEM from unmethylated probes (95% on average) and a majority of hyperSEM from methylated probes (52% on average); the total number of SEM also decreased, more significantly than in GSE55763 (from ∼1,600,000 to ∼900,000) (**Figure 5c**).

**FIGURE 5.**
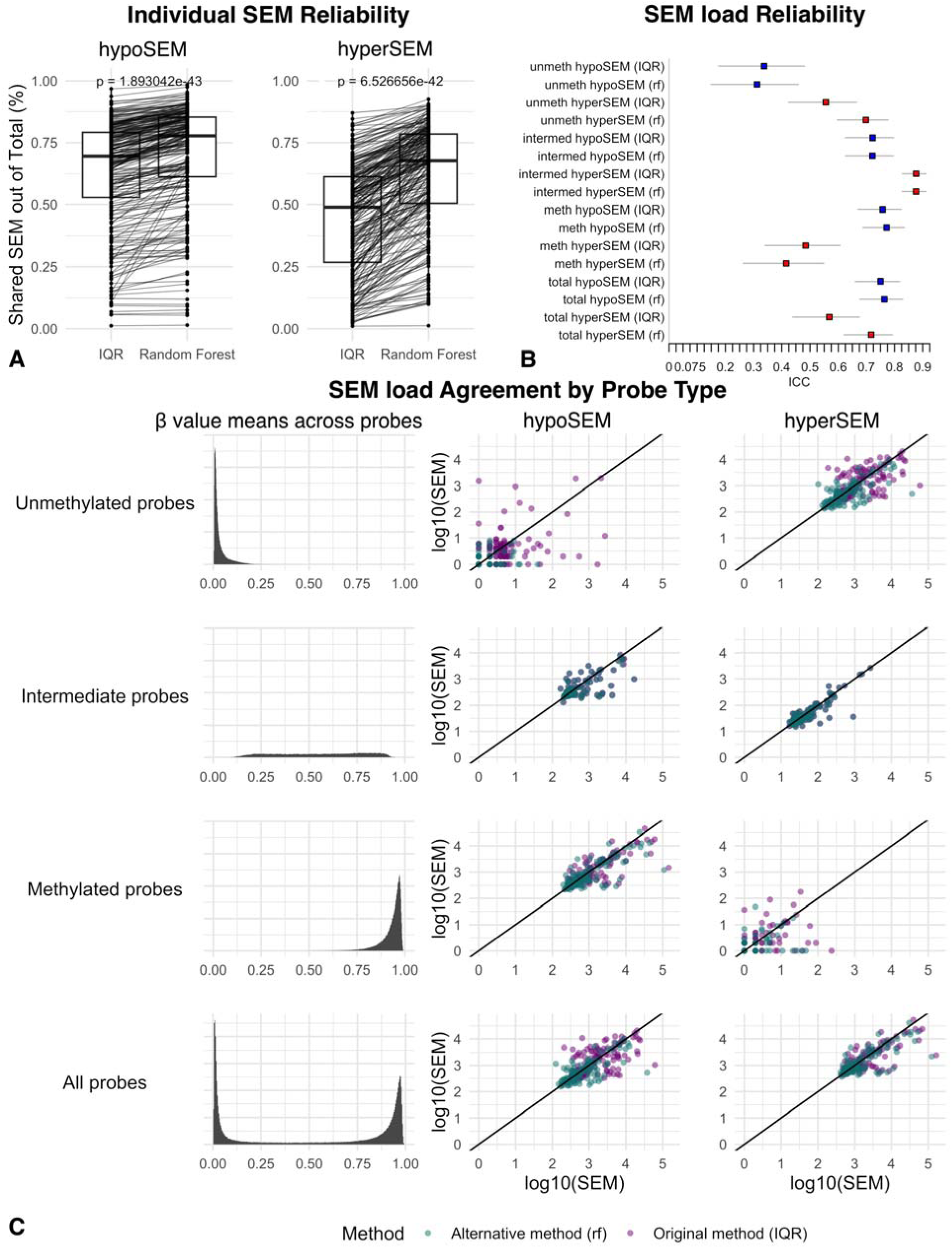
SEM reliability: Effect of random forest filter in the Sister Study. Figure 5a: Boxplots illustrating the proportion of shared hypoSEMs (left) and hyperSEMs (right) out of the total number of identified hypo- and hyperSEMs, respectively, as detected by the original IQR-based method vs filtered by the RF-based method within the Sister Study. The different boxplots represent the results obtained by the two SEM detection methods, with lines connecting the same samples analyzed by the two methods. The replicate batch representation is omitted from this representation for clarity. Figure 5b: Forest plot of the ICC scores for SEM load subtypes within the Sister Study. The hypoSEM loads are color-coded in blue, while the hyperSEM loads are depicted in red. Horizontal bars mark the lower and upper bounds of the ICC measurements. The label ‘Total’ signifies the SEM loads derived from all probes, while labels ‘unmeth,’ ‘intermed,’ and ‘meth’ specifically denote the SEM loads originating from unmethylated, intermediate, and methylated probes, respectively. The tag (rf) indicates that the SEMs were detected with the RF-based filtering, while the rest were detected using the IQR-based method. Figure 5c: Scatterplots for the SEM subtypes showcasing the agreement in SEM loads between the first and second batches of replicates within the Sister Study. SEM loads detected with the original method are illustrated in green, while those identified with the RF-based filtering are depicted in purple.

### Impact of Reference Dataset Size on SEM Reliability

We utilized GSE55763 to assess the impact of reference dataset size on SEM reliability. We found that ICC scores plateaued when the reference dataset reached approximately 50 samples and the proportion of shared SEMs plateaued at 150 samples (**Supplementary Figure 5**).

### Association between SEMs and Cardiovascular Aging Outcomes

Lastly, we extended our analyses to the Framingham Heart Study (FHS) dataset to test SEM associations with mortality and age-related phenotypic data. We observed the following correlations with age: hypoSEMs (r = 0.24; p = 1.13e-50) and hyperSEMs (r = 0.18; p = 1.58e-28) detected with the original method; and hypoSEMs (r = 0.23; p = 3.39e-50) and hyperSEMs (r = 0.20; p = 8.69e-37) detected with the RF method (**Figure 6a**).

**FIGURE 6.**
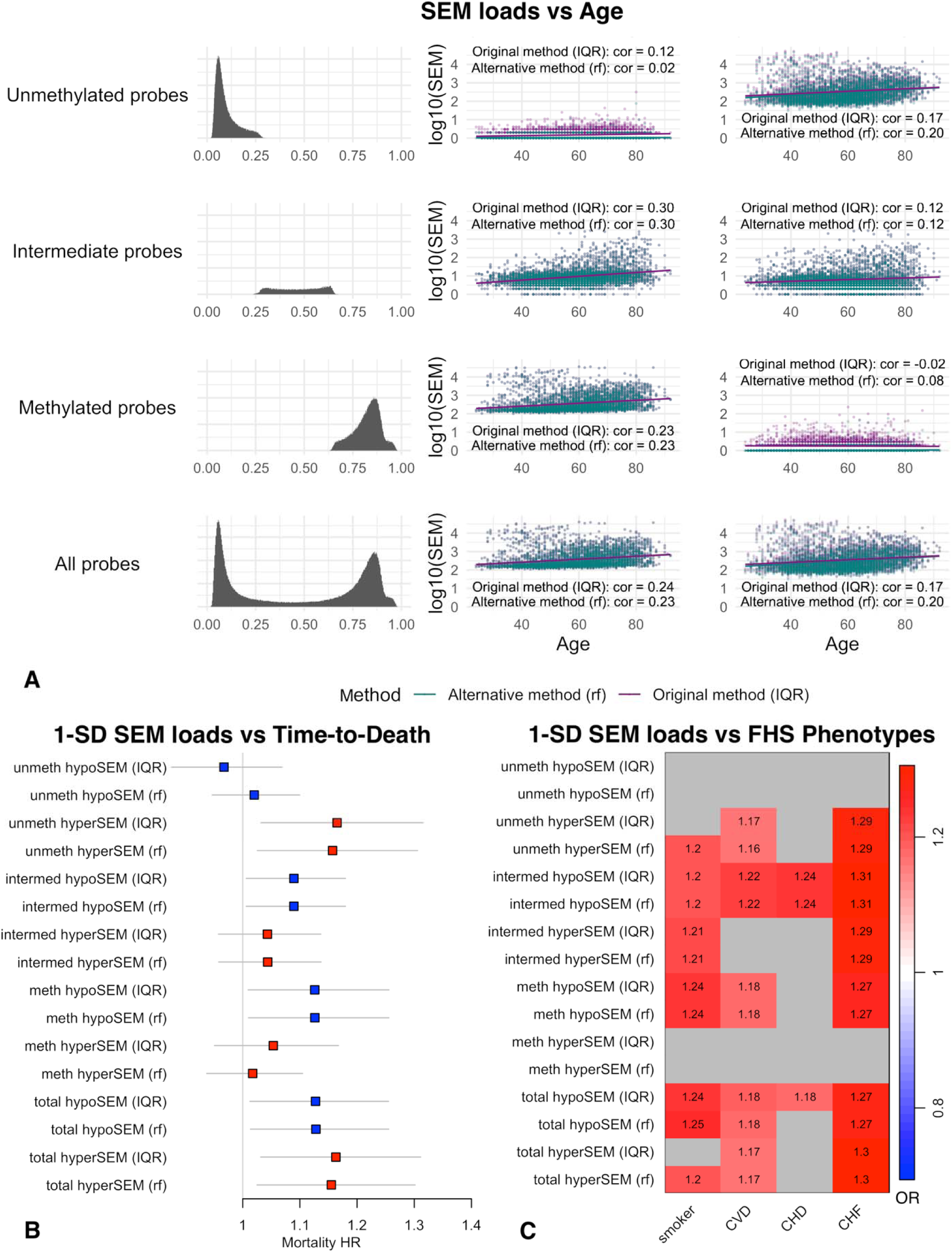
SEM associations with age, mortality, and cardiovascular phenotypes. Figure 6a: Scatterplots illustrating the association between SEM subtype loads (log10) and age within the Framingham Heart Study (FHS) dataset. SEM loads detected by the conventional method are color-coded in green, while those identified utilizing the RF-based filtering method are portrayed in purple. Figure 6b: Forest plot of the Hazard Ratios (HR) of standardized SEM load subtypes in relation to mortality within the FHS dataset adjusted for age and sex. The hypoSEM loads are represented in blue, while the hyperSEM loads are in red. Horizontal bars indicate the 95% confidence interval for each HR. The label ‘total’ refers to the SEM loads extracted from all probes, whereas labels ‘unmeth,’ ‘intermed,’ and ‘meth’ explicitly represent the SEM loads originating from unmethylated, intermediate, and methylated probes, respectively, as categorized through the reference dataset analysis. The annotation (rf) specifies that the SEMs were filtered using the RF-based filtering method. Figure 6c: Heatmap of associations between standardized SEM load subtypes and phenotypic traits within the Framingham Heart Study (FHS) dataset, with adjustments made for age and sex. The color corresponds to odds ratios (OR) of respective phenotype in relationship to 1 standard deviation increase in SEM load subtypes. Associations that did not reach statistical significance (Bonferroni corrected) are colored in gray. The label ‘Total’ signifies the SEM loads derived from all probes. Labels ‘unmeth,’ ‘intermed,’ and ‘meth’ indicate the SEM loads originating from unmethylated, intermediate, and methylated probes, respectively. The annotation (rf) indicates that the SEMs were identified using the RF-based filtering method, contrasting the rest which were detected utilizing the IQR-based method.

Upon examining standardized total SEM loads for potential associations with time-to-death while controlling for age and sex, we found statistically significant associations for both IQR (HR 1.149, CI 95% 1.023–1.290; p = 0.0187) and RF detection methods (HR 1.144, CI 95% 1.020–1.283; p = 0.022). Next, we analyzed associations with mortality of all six SEM subtypes and found that only three of them had statistically significant associations with mortality; one standard deviation (sd) increase in hyperSEMs from unmethylated probes (HR 1.165, CI 95% 1.032–1.315; p = 0.0135), hypoSEMs from intermediate probes (HR 1.090, CI 95% 1.006–1.180; p = 0.0347), and hypoSEMs from methylated probes (HR 1.126, CI 95% 1.010–1.256; p = 0.0331) was associated with increased risks of mortality. The RF-based detection method maintained these associations (**Figure 6b**). Combining hypoSEMs from both intermediate and methylated probes only marginally enhanced the associations with mortality (HR 1.129, CI 95% 1.012-1.263; p = 0.0287). When comparing the performance of the Cox proportional hazards models with hypoSEMs from methylated probes against hypoSEMs from both methylated and intermediate probes, minimal differences were observed in both AIC and BIC (ΔAIC = 0.228; ΔBIC = 0.228). Additionally, we conducted a sensitivity analysis and determined that adjusting for inferred cell type composition strengthens the associations with mortality for hyperSEMs from unmethylated probes (HR 1.236, CI 95% 1.097-1.391; p = 4.7e-4) and hypoSEMs from methylated probes (HR 1.172, CI 95% 1.053-1.304; p = 3.60e-3). The mortality for hypoSEMs from intermediate probes was slightly reduced (HR 1.084, CI 95% 0.999-1.177; p = 0.052) (**Supplementary Figure 6**).

We evaluated the associations between SEM loads and available phenotypic data, controlling for age and sex, and found multiple associations with cardiovascular diseases including coronary heart disease and congestive heart failure (**Figure 6c**). These associations remained after adjusting for sex and smoking status, with congestive heart failure exhibiting the strongest associations: a 1-SD increase in SEM loads adjusted for age and sex had increased odds ratio for hyperSEMs from unmethylated probes (OR 1.29; p = 1.98e-4), hypoSEMs (OR 1.30; p = 6.13e-5) and hyperSEMs from intermediate probes (OR 1.28; p = 2.03e-4), and for hypoSEMs from methylated probes (OR 1.26; p = 4.97e-4). The RF-based method preserved these associations (**Figure 6c**). Finally, after adjusting for inferred cell counts, the odds ratios increased to OR 1.33 (p = 2.81e-5) for hyperSEMs from unmethylated probes, OR 1.36 (p = 5.26e-6) for hypoSEMs and hyperSEMs (p = 6.14e-6) from intermediate probes, and OR 1.34 (p = 1.35e-5) for hypoSEMs from methylated probes.

### SEMdetectR: a new tool for SEM analysis

To meet the need for robust tools in SEM analysis, we have developed a new R package: *SEMdetectR* (**Figure 7**). Engineered to utilize parallel programming, this tool facilitates expedited SEM detection using either IQR- or RF-based techniques, bolstered by a spectrum of filtering and analytical capabilities. *SEMdetectR* has the flexibility to detect SEM in all probes or any subset of probes specified by user, employ reference methylation data derived from our whole-blood dataset investigations, or cluster their reference DNAme data based on methylation status. Our additional analyses revealed a significant overlap in terms of unmethylated (161,754 probes) and methylated (187,689 probes) probes across the three datasets we investigated, and these probes are included with the R package. By offering a standardized approach, we anticipate that more researchers will delve into SEM studies, propelling us closer to unraveling how epigenetics shapes aging, health, and disease.

**FIGURE 7.**
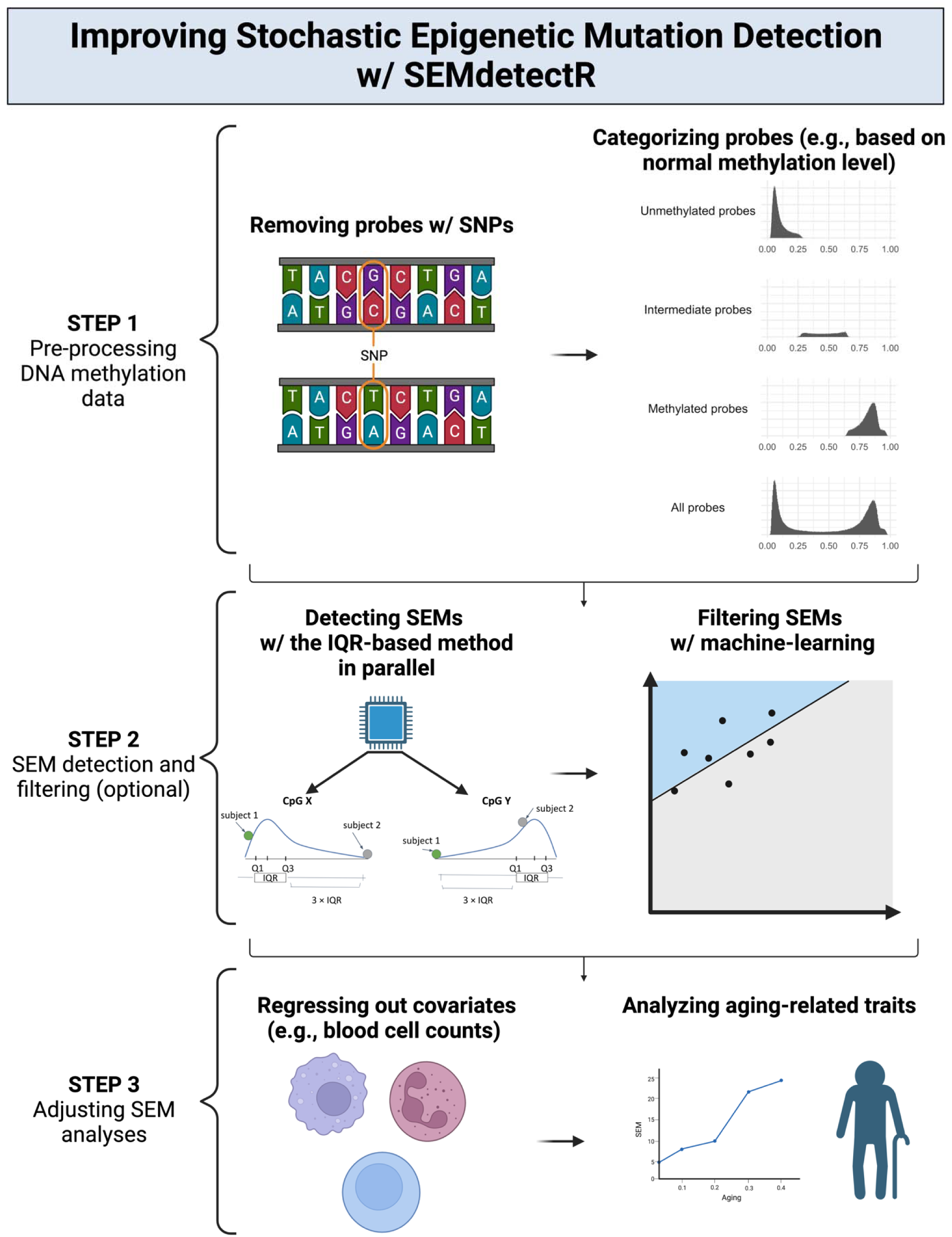
Overview of the Improved Workflow for Stochastic Epigenetic Mutation Detection Using *SEMdetectR*. Illustration of a step-by-step *SEMdetectR* pipeline, highlighting optional actions at each step for improved SEM analysis. **Step 1:** The workflow begins with the pre-processing of DNA methylation data, where probes containing SNPs can be removed to mitigate genetic influences. Probes can then be categorized based on their methylation levels into unmethylated, intermediate, or methylated groups. Users can specify custom probe groupings. **Step 2:** SEMs are identified using the IQR-based method, which can be applied in parallel to enhance computational efficiency. An optional filtering step can be employed, utilizing the random forest model to refine the selection of SEMs. **Step 3:** Adjustments for covariates such as blood cell counts can be incorporated to correct for their potential confounding effects on subsequent SEM analyses.

## DISCUSSION

In this study, we examined the reliability and biological relevance of stochastic epigenetic mutations (SEMs) across multiple datasets. We describe the reliability of both individual SEMs and SEM load, uncover novel associations with mortality and age-associated phenotypes related to cardiovascular health, and find that specific subsets of SEMs are both reliable and associated with aging outcomes.

The GSE55763 dataset was particularly useful for technical reliability analysis, given its inclusion of 36 pairs of technical replicates and 2,664 samples measured once, acting as a reference dataset for SEM detection and allowing for in-depth examination of how various probe and sample characteristics impact SEM reliability.

While the SEM intraclass correlation coefficient values (∼0.96 for hypoSEM and ∼0.90 for hyperSEM) may be interpreted as markers of high reliability <u>[24]</u>, our earlier work suggests that similarly high ICCs for epigenetic clocks are insufficient for multiple use cases, particularly in longitudinal or interventional contexts, and that strategies to bolster ICCs to >0.99 are beneficial <u>[22]</u>. This raises questions about the adequacy of the current SEM detection strategy and underscores the need for its refinement. More crucially, despite relatively high ICC scores for SEM load between technical replicates, the low reliability at the level of individual SEMs (many unshared SEMs between replicates) demonstrates that a significant proportion of SEMs represent technical noise, which is particularly problematic for downstream analyses of individual SEMs. For example, it has been suggested that the different SEM patterns between people with the same environmental exposure can help explain divergent health outcomes resulting from that exposure <u>[10]</u>. However, if many individual SEMs are noise, then this hypothesis is much more difficult to test. This illustrates a need for development of SEM detection strategies that can discern genuine biological signals from technical artifacts.

Our findings underscore that there is not a single overarching factor, but rather a complex interplay of multiple determinants shaping SEM reliability, including probe and sample characteristics such as variance and cell composition. In particular, the lower IQR and standard deviation in unshared SEMs compared to shared SEMs aligns with prior findings of reduced reliability for CpGs with low variance <u>[20]</u>. We find that integrating these determinants into a predictive model can effectively filter out unreliable SEMs. Our random forest model increased the proportion of shared SEM and concurrently increased the ICC scores in both the original GSE55763 dataset as well as the Sister Study test dataset. It is important to note that the RF-based method is computationally more intensive than the original method. The translatability of the model across datasets is consistent with prior findings that CpG beta-value ICCs are similar across different technical replicate datasets <u>[19,25]</u>.

SEMs, especially hyperSEMs, are sensitive to variations in cell type composition, consistent with prior observations <u>[6]</u>. Because a change in cell composition would be present in both technical replicates, such SEMs are more likely to be reliable. Blood cell composition is known to change with aging and may contribute to mortality associations for epigenetic clocks given that adjusting for cell counts can reduce epigenetic clock mortality associations <u>[26]</u>. However, we find that adjusting for cell composition can enhance the associations between SEM load and aging outcomes. Interestingly, the associations with mortality were independent of smoking status, which is again different than epigenetic clocks which are known to predict mortality at least partially via smoking <u>[27]</u>. <u>These findings support examining SEM loads both before and after adjustment for cell composition, as well as other potential confounders or mediating variables.</u>

At the same time, the presence of SNPs, especially at the Single Base Extension site (SBE), was specifically associated with increased hypoSEM reliability. A major concern is that these SNP-associated SEMs may reflect genetic, rather than epigenetic, variability, since such SNPs would be present in both technical replicates and have a large effect on measured methylation and might have limited relevance for aging studies. However, this is likely a minor issue, as others have reported SEM load is not heritable and most SEMs are not shared between tissues from the same person <u>[6,8,10]</u>, and we found only a small proportion of SEMs are SNP-associated and removing these probes has minimal impact on ICC scores. <u>Regardless, our advice leans towards omitting probes with SNPs in aging-focused analyses.</u>

Our results indicate that the utility of defining multiple SEM subtypes should be further investigated. As expected, hypo- and hyper-SEM differ in reliability and associated probe and sample characteristics, supporting the notion that they stem from distinct underlying factors and represent separate biological phenomena. We find that hypoSEMs predominantly come from methylated probes and hyperSEMs mainly originate from unmethylated ones. We define six SEM subtypes, based on their baseline methylation status: hypo- and hyperSEMs from unmethylated, intermediate, and methylated probes respectively. HypoSEMs from unmethylated probes and hyperSEMs from methylated ones have markedly reduced reliability as well as null associations with all-cause mortality and cardiovascular disease, hinting that these specific SEM subcategories might contain unwanted noise in SEM data. <u>In contrast, hypoSEMs from methylated probes and hyperSEMs from unmethylated probes show both good reliability and associations with mortality and cardiovascular disease, indicating that future studies should pay particular attention to these SEM subtypes</u>. It is notable that prior studies primarily utilized total SEM load when examining risk for mortality and age-related diseases such as cancer <u>[8,9]</u>. Our results further suggest that it is important to examine individual SEM subtypes as some subsets display stronger associations and reliability than others, while simply utilizing total SEM loads can obscure which SEM subtypes are most important.

It should be noted that there is considerable potential to further delineate SEM subtypes beyond the categorization based on probe methylation status. For example, SEMs have previously been reported to be enriched in specific biological pathways or genomic locations <u>[9,10,13]</u>. One could define SEM subtypes based on genomic location or regulatory features, which we found can vary in SEM reliability. It would be interesting to determine if SEM subtypes based on biological pathways predict different phenotypic consequences, or if some biological pathways are more vulnerable to unreliable SEMs. Furthermore, future research could delve into alternative methods for computing SEM loads, such as using factor analysis, in datasets abundant with phenotypic details associated with aging. We anticipate that *SEMdetectR* will serve as a valuable asset for such research directions.

Notably, we discover associations between SEMs and cardiovascular diseases (CVD) including coronary heart disease and congestive heart failure. Previous research has corroborated the idea that alterations in DNA methylation play a role in controlling the biological mechanisms behind CVD, such as the progression of atherosclerosis, the management of blood pressure in hypertension, and the inflammatory responses <u>[28]</u>. Thus, our findings contribute to the growing body of research on the influence of epigenetics in cardiovascular health and underscores the potential for SEMs to serve as biomarkers for cardiovascular risk assessment.

SEMs must always be defined relative to a reference population. Our analyses of the impact of sample size on SEM reliability in GSE55763 highlight that after a specific threshold (approximately 50 samples for ICC scores and 150 samples for the proportion of shared SEMs), increasing the reference dataset size does not significantly enhance the reliability. <u>Thus, we generally recommend at least 150 samples for a reference dataset, with a wide age range and inclusion of both sexes.</u> Interestingly, ICC scores in the Sister Study were lower in general than GSE55763, and hypoSEMs in particular did not improve in ICC in the Sister Study with the random forest model (though the proportion of shared hypoSEMs did). This may reflect the study’s exclusively female demography and corresponding reduced biological variation. The impact of reference and study population characteristics should be a focus of future studies.

Our analyses, investigating technical reliability and aging associations of SEMs and introducing a novel software tool with the potential to enable consistent interpretations of SEM studies, advance the understanding and methodological framework for SEM analysis. Additionally, the associations discovered between SEMs and critical health outcomes highlight the potential utility of SEM analysis in aging research and possibly in the broader field of epigenetics and age-related diseases. Future studies can utilize the insights and best practices identified here to investigate SEMs in a variety of studies concerning diverse populations, longitudinal or interventional datasets, multiple tissue types, various diseases, or other methylation measurement platforms.

## METHODS

### Datasets

#### GSE55763

Bisulfite-treated DNA samples from 2,664 human subjects’ peripheral blood, along with 36 technical duplicates, from the London Life Sciences Prospective Population (LOLIPOP) study were analyzed using Illumina Infinium HumanMethylation450 BeadChip <u>[29]</u>. The dataset underwent quantile normalization, which, according to Lehne et al., demonstrated the highest consistency between technical replicates among 10 normalization techniques. Additionally, control probes were employed to mitigate systematic technical discrepancies, such as those arising from batches and plates. The dataset contains both males (n=27) and females (n=9) with age ranging from 37 to 74 (mean 53).

#### NIEHS Sister Study (GSE174422)

In the context of the existing National Institute of Environmental Health Science (NIEHS) Sister Study, DNA from blood samples of 128 technical duplicate pairs from women were assayed using Illumina Infinium HumanMethylation450 BeadChip <u>[21]</u>. Genomic DNA was extracted from aliquots of whole blood and was randomly allocated across both plates and arrays, ensuring duplicates of any given sample were bisulfite-converted on separate plates and analyzed on different arrays. The age range is 36.6 to 75.1 (mean 57.6).

#### Framingham Heart Study

The Framingham Heart Study (FHS) dataset was previously described <u>[25,30,31]</u> and encompasses 2,748 participants from the Offspring cohort who attended the eighth examination phase (2005-2008) and 1,457 from the Third Generation group present during the second examination cycle (2005-2008). Illumina Infinium HumanMethylation450 BeadChip was employed to assay DNA methylation. The study was approved by the IRB at Boston University Medical Center, and all participants provided written informed consent at the time of each examination visit.

### SEM Detection and Analyses

#### SEM detection with IQR-based method

To evaluate SEM reliability, both for individual SEMs and cumulative SEM loads, we used the interquartile range (IQR)-based method for outlier detection, as detailed by <u>[7]</u>. For every locus, this method identifies hypoSEMs as DNA methylation outliers that fall 3 x IQR below the first quartile (Q1) and hyperSEMs as those lying 3 x IQR above the third quartile (Q3). IQR was calculated based on 2,664 samples without replicates in the case of the GSE55763 dataset; based on all technical replicates in the case of the Sister Study dataset; and based on all samples in the case of the FHS dataset. CpGs with IQR = 0 were removed, since SEMs are undefined for these CpGs. CpGs were processed in parallel with *foreach* (version 1.5.2) and *doParallel* (version 1.0.17).

#### SEM loads

SEM loads were defined as the cumulative count of SEMs for each individual, separated into hypoSEM and hyperSEM loads (or further subtypes based on methylation status). We analyzed these by taking the log10 of the SEM load, consistent with earlier aging studies <u>[6,7]</u>.

#### SEM associations and statistical differences between groups

The statistical difference in SEM loads between replicates and detection methods, as well as SEM reliability status across continuous features, were assessed with Wilcoxon tests (two-sample for paired measurements) using R package *stats* (version 4.2.3). Differences in SEM reliability status across categorical features were assessed with Chi-squared tests using R package *stats* (version 4.2.3). Pearson correlations between SEM loads and age were calculated using R package *stats* (version 4.2.3). The associations between SEM loads and FHS phenotypic variables were assessed with logistic regressions using *stats* (version 4.2.3), and the association with mortality with Cox proportional hazard regression models using R package *survival* (version 3.5.7). Models were standardized, meaning that they represent the effect of a 1 standard deviation (SD) change in SEM load; for continuous phenotypes, they reflect a 1-SD change in the phenotype. Expectations-Maximization analysis of deltaIQR distributions was performed with R *mixtools* package (version 2.0.0).

#### ICC scores

The ICC scores were computed using R package *irr* (version 0.84.1), employing a single-rater, absolute-agreement, two-way random-effects model, as previously described <u>[25]</u>.

#### Visualization

Visual representations were created utilizing R packages *ggplot2* (version 3.4.3), *forestplot* (version 3.1.3), and *ComplexHeatmap* (version 2.13.0).

### Probe and Sample Characterization

#### Probe statistics

Mean, median, quartiles, sum, standard deviation, median absolute deviation, minimum, and maximum of probes were calculated using R package *stats* (version 4.2.3); skewness and kurtosis were calculated using R package *moments* (version 0.14.1). IQR, coefficient of variation, and range were derived from these statistics.

#### Probe annotations

Probe annotations were retrieved using R package *minfi* (version 1.44.0).

#### Probe clustering

Probe clustering based on methylation status was done using probe summary statistics and kmeans from R package *stats* (version 4.2.3).

#### Cell type composition inference

We utilized the method described by <u>[32]</u> to estimate white blood cell sub-populations. The code was downloaded from the data repository associated with the publication.

### Random Forest Training

#### Feature selection

We prioritized features most strongly associated with ground truth labels (above 0.1 in association strength) and selected only one feature from each set of associated features. For example, “mean” was chosen over “median” from the mean/median pair, and SNPs at SBE were preferred over SNPs at CpG sites. The strengths of associations were assessed with cramer’s v with *confintr* package (version 1.0.2) (categorical features), point-biserial correlation (numerical vs binary features, and numerical vs ordinal categorical after one-hot encoding) and Pearson correlation coefficients with R package *stats* (version 4.2.3) (numerical features).

#### Model training

We performed a random 80/20 split for training and testing data, used 5-fold cross-validation and tuned the number of features considered in individual decision trees, sample size, and number of trees using *mlr* (version 2.19.1). Gini coefficients were analyzed using *randomForest* (version 4.7-1.1).

## CODE/DATA AVAILABILITY

The datasets comprising technical replicates utilized in this research are publicly accessible on the NCBI Gene Expression Omnibus (GEO) under accession numbers GSE55763 and GSE174422. However, due to the sensitive nature of the health data contained within the Framingham Heart Study (FHS) dataset, researchers interested in accessing this data will need to submit an application through the database of Genotypes and Phenotypes (dbGaP) at https://dbgap.ncbi.nlm.nih.gov/aa/ (dbGaP accession number: phs000724.v7.p11). The SEMdetectR software package developed as part of this study will be made available on GitHub upon publication. The repository will include the source code, alongside comprehensive documentation to facilitate utilization by other researchers in the community.

## AUTHOR CONTRIBUTION

YM, MEL, and AHC conceived the project and study design and obtained and cleaned the data. YM performed all analyses, trained the machine learning models, generated the figures, and developed the SEMdetectR software package. ML and AHC supervised, provided feedback, and provided code for analysis of mortality and reliability. YM and AHC authored the manuscript. All authors reviewed and approved the manuscript.

## ACKNOWLEDGEMENTS

This project was principally supported by the funding awarded to YM from the Biomarker Network (R24 AG037898), and to AHC and MEL by the National Institute on Aging (R01AG057912 and R01AG065403).

