## Supplementary Figures for "Stochastic Epigenetic Mutations: Reliable Detection and Associations with Cardiovascular Aging"

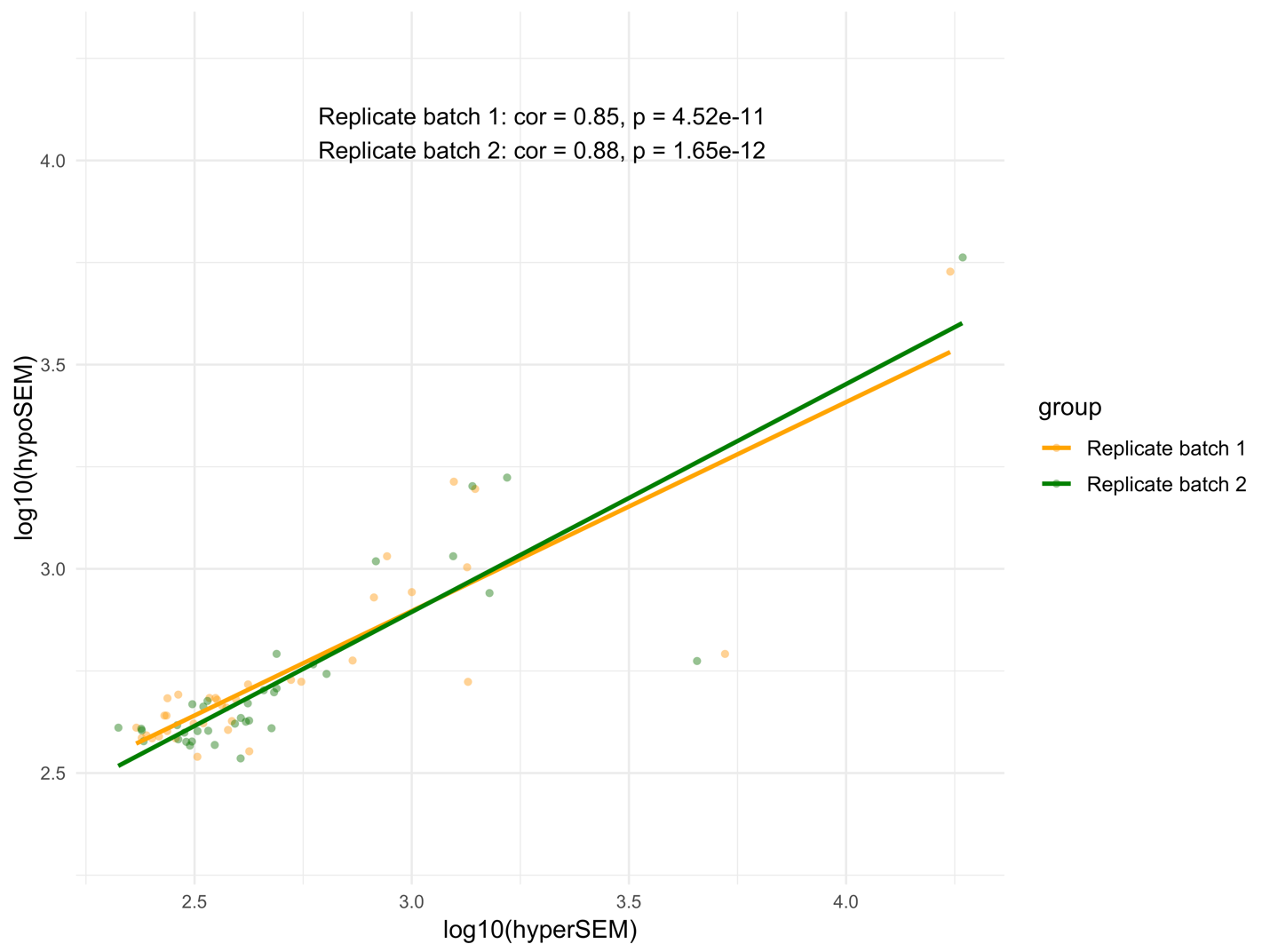

**SUPPLEMENTARY FIGURE 1. HypoSEM vs hyperSEM loads in GSE55763**

The two batches of replicates are distinguished by color, with one batch represented in yellow and the other in green.

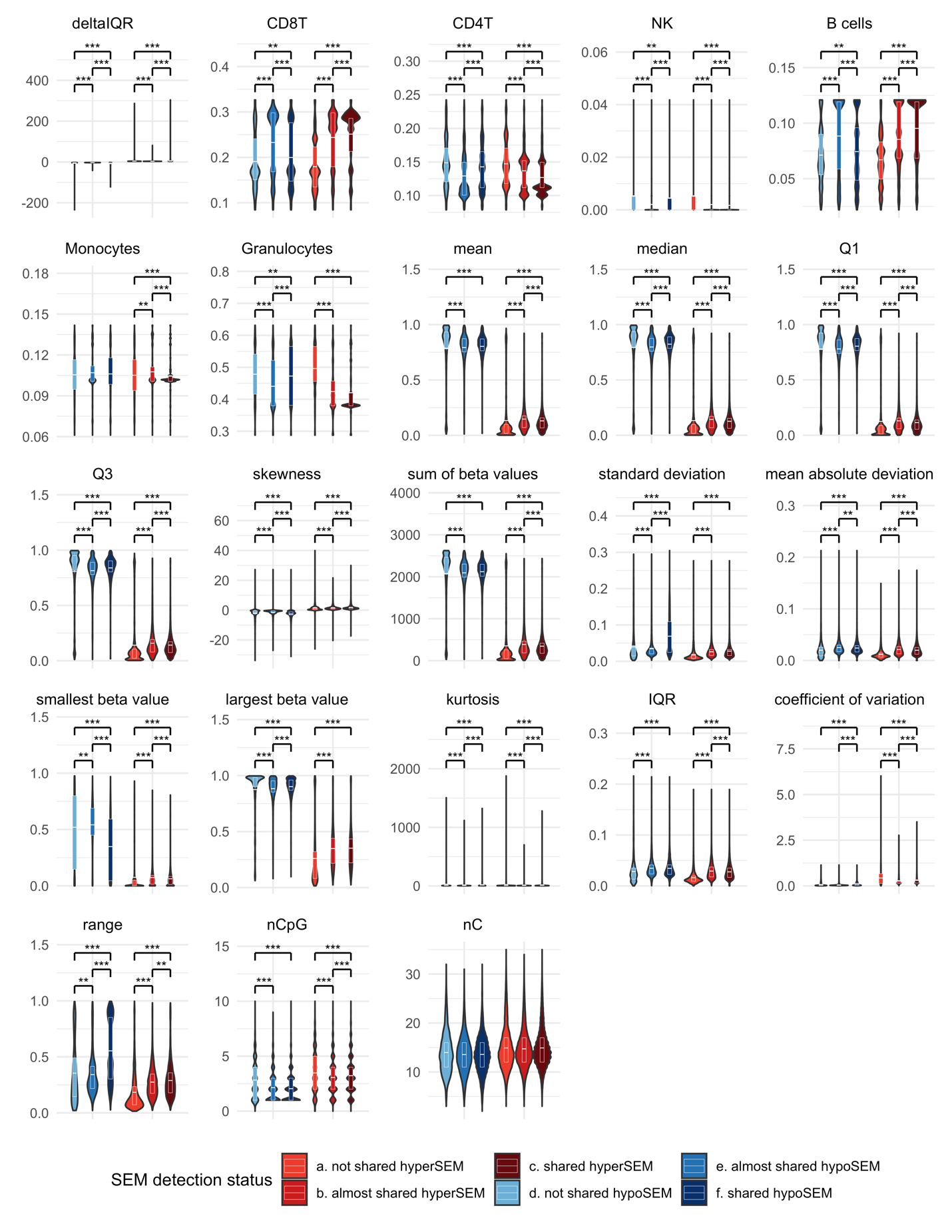

**SUPPLEMENTARY FIGURE 2. Numerical features vs SEM reliability status in GSE55763**

HypoSEMs are depicted in blue and hyperSEMs in red. Statistically significant differences between violin plots are marked by asterisks, with white horizontal lines signifying the mean and quartile values. The significance values are annotated as follows: *** indicates p < 2e-16 and ** indicates p < 3.8e-4 (Bonferroni corrected significance value).

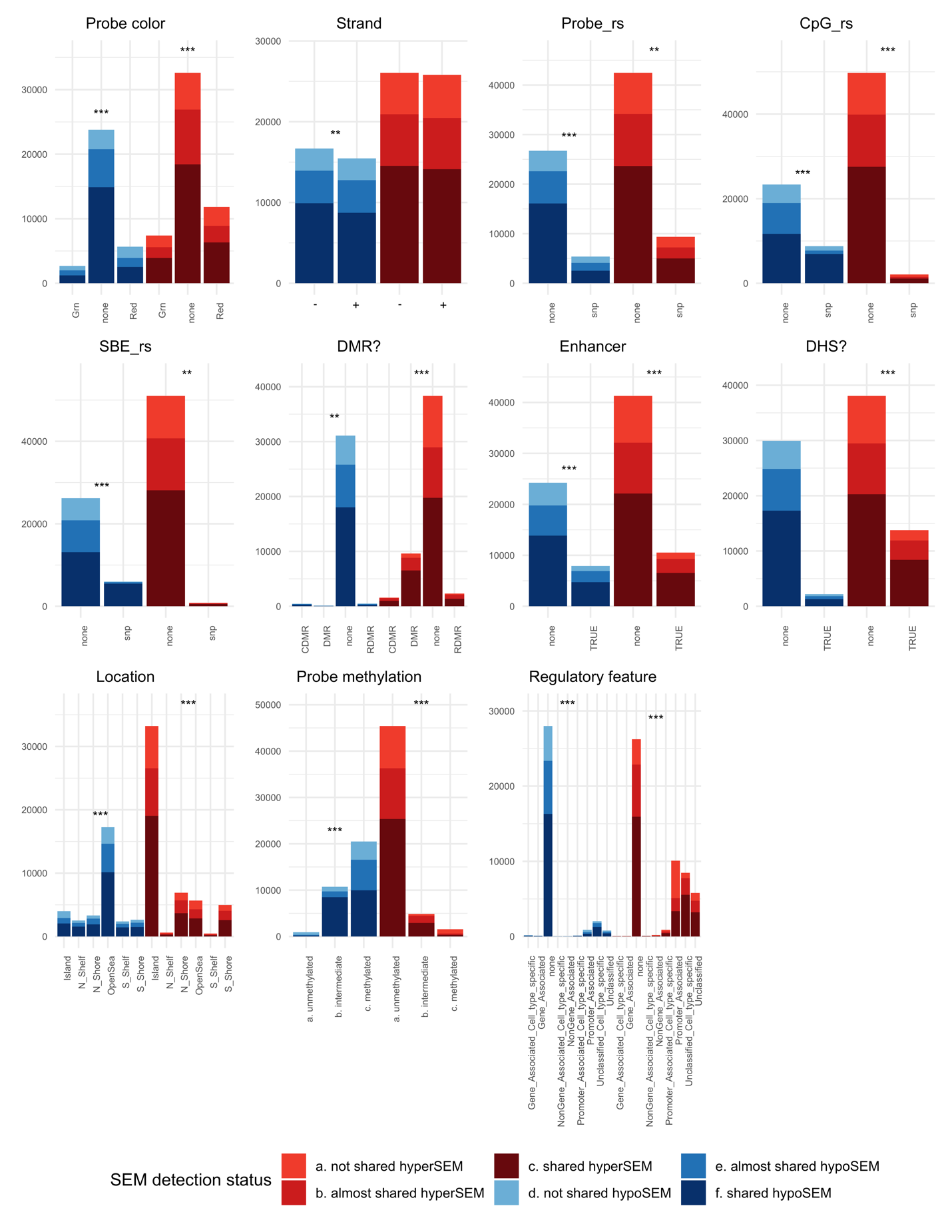

**SUPPLEMENTARY FIGURE 3. Categorical features vs SEM reliability in GSE55763**

HypoSEMs are represented in blue and hyperSEMs in red. The features are depicted through stacked bar plots. Asterisks positioned atop the bar plots denote statistically significant differences within the hypo- or hyperSEM groups. Significance values are annotated as follows: *** indicates p < 2e-16, and ** indicates p < 3.8e-4 (Bonferroni corrected significance value.

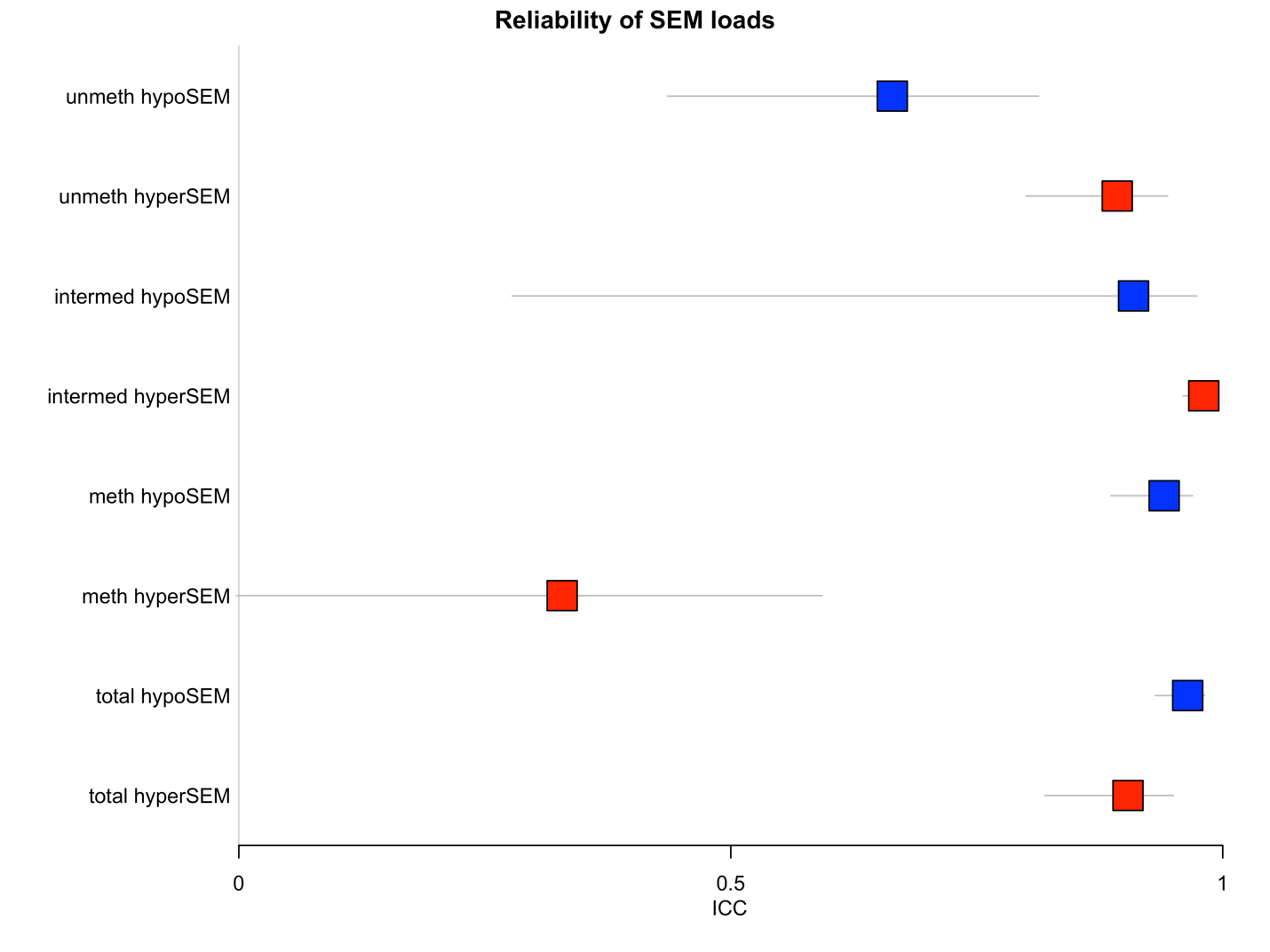

**SUPPLEMENTARY FIGURE 4. Intraclass Correlation Coefficient (ICC) scores for SEM loads in GSE55763**

The hypoSEM loads are depicted in blue, while the hyperSEM loads are in red. The horizontal bars indicate the lower and upper bounds of the ICC measurements. The label 'total' refers to the SEM loads derived from all DNA methylation probes, whereas 'unmeth', 'intermed', and 'meth' specifically denote the SEM loads originating from unmethylated, intermediate, and methylated probes, respectively, as determined through the reference dataset analysis.

|  | **ICC** | **LowerBound** | **UpperBound** |
| --- | --- | --- | --- |
| **unmeth_hypo** | 0.6641548 | 0.435710659 | 0.8127934 |
| **unmeth_hyper** | 0.8927819 | 0.799933826 | 0.9438685 |
| **mid_hypo** | 0.9093573 | 0.277849017 | 0.9738080 |
| **mid_hyper** | 0.9806915 | 0.959925890 | 0.9904284 |
| **meth_hypo** | 0.9404630 | 0.886399800 | 0.9692044 |
| **meth_hyper** | 0.3286415 | -0.002323169 | 0.5925104 |
| **sum_hypo** | 0.9643790 | 0.931300282 | 0.9816788 |
| **sum_hyper** | 0.9036928 | 0.819258851 | 0.9497387 |

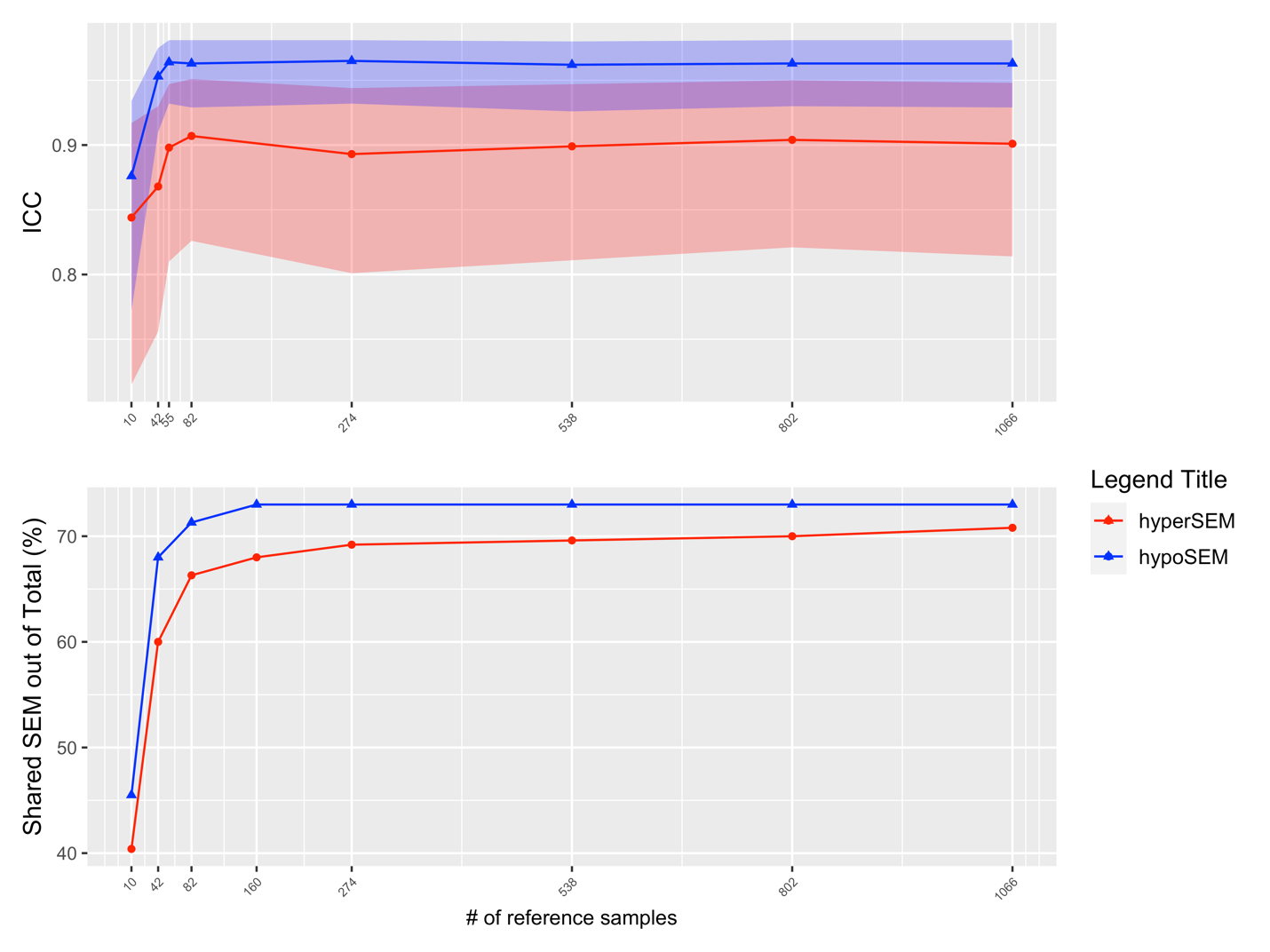

**SUPPLEMENTARY FIGURE 5. Effect of reference dataset size on technical reliability of SEMs**

Intraclass correlation coefficients (top) and the percentage of shared SEMs (bottom) are compared for different numbers of reference samples. HypoSEM values are in blue and hyperSEM values are in red. The x-axis is non-uniformly scaled in both scenarios to accentuate the sharp enhancement of reliability with minor increments in the reference dataset size.

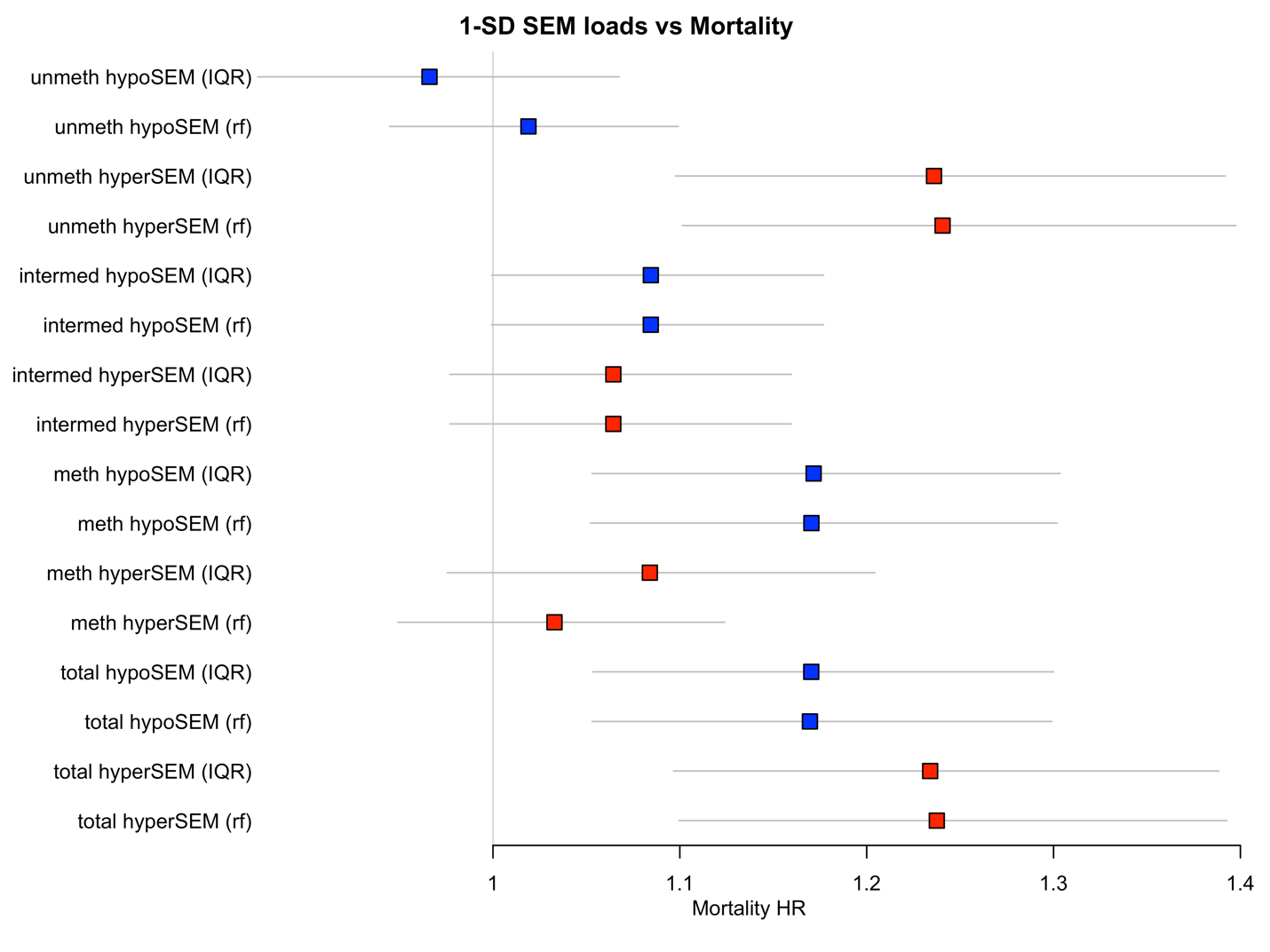

**SUPPLEMENTARY FIGURE 6. Mortality associations of SEM loads adjusted for age, sex, and inferred cell type composition**

The forest plot is shown with the standardized hazard ratios (HR) of different subtypes of SEM, indicating the effect of a 1-SD change in SEM load. The hypoSEM loads are represented in blue, while the hyperSEM loads are in red. Horizontal bars indicate the 95% confidence interval for each HR. The label 'total' refers to the SEM loads extracted from all probes, whereas labels 'unmeth,' 'intermed,' and 'meth' represent the SEM loads originating from unmethylated, intermediate, and methylated probes, respectively. The annotation (rf) specifies that the SEMs were filtered using the RF (random forest) filtering method.

| **Clock** | **MortalityHR_SD** | **MortalityHR_SDUpper** | **MortalityHR_SDLower** | **MortalityP** |
| --- | --- | --- | --- | --- |
| unmeth hypoSEM | 0.9660385 | 1.067644 | 0.8741029 | 0.4983031763 |
| unmeth hypoSEM (rf) | 1.0189929 | 1.099185 | 0.9446513 | 0.6264080790 |
| unmeth hyperSEM | 1.2360346 | 1.391920 | 1.0976077 | 0.0004708898 |
| unmeth hyperSEM (rf) | 1.2406123 | 1.397599 | 1.1012593 | 0.0003902584 |
| intermed hypoSEM | 1.0844789 | 1.176950 | 0.9992728 | 0.0520719309 |
| intermed hypoSEM (rf) | 1.0844503 | 1.176914 | 0.9992510 | 0.0521352238 |
| intermed hyperSEM | 1.0644292 | 1.159729 | 0.9769607 | 0.1535266570 |
| intermed hyperSEM (rf) | 1.0644019 | 1.159705 | 0.9769305 | 0.1537210862 |
| meth hypoSEM | 1.1716090 | 1.303589 | 1.0529910 | 0.0036368776 |
| meth hypoSEM (rf) | 1.1704251 | 1.302035 | 1.0521187 | 0.0037984649 |
| meth hyperSEM | 1.0839499 | 1.204400 | 0.9755460 | 0.1337582952 |
| meth hyperSEM (rf) | 1.0329260 | 1.124137 | 0.9491156 | 0.4530490622 |
| total hypoSEM | 1.1703314 | 1.300085 | 1.0535273 | 0.0033681543 |
| total hypoSEM (rf) | 1.1696279 | 1.299057 | 1.0530945 | 0.0034328067 |
| total hyperSEM | 1.2339627 | 1.388458 | 1.0966579 | 0.0004776521 |
| total hyperSEM (rf) | 1.2375237 | 1.392691 | 1.0996441 | 0.0004062375 |
